# Participant-specific macromolecule information improves quantification of GABA-edited spectra at 3T

**DOI:** 10.64898/2026.10.07.757300

**Authors:** Justin R. Singer, Kimberly L. Chan

**Author notes:** Corresponding Author: Kimberly L. Chan, Ph.D. Advanced Imaging Research Center, UT Southwestern, Medical Center, 5323 Harry Hines Boulevard, NE.3.102, Dallas, TX 75390.

## Abstract

**Purpose:** To evaluate the effects of different MM fitting strategies on gamma-aminobutyric acid (GABA) estimates using MEGA-PRESS at 3T.

**Methods:** Thirteen healthy individuals were scanned in the parietal lobe and centrum semiovale. GABA-edited scans were performed at TE=64ms (GABA+MMs) and TE=80ms (MM-suppressed GABA). MM-edited spectra were acquired using MEGA-PRESS with an inversion pre-pulse. MM peaks at 3.0 (MM_3.0_), 2.05 (MM_2.05_), and 0.94 (MM_0.94_) ppm were parameterized to form five MM basis categories. Cohort-mean/individual MM basis sets were constructed with all MM peaks combined (MM_together_), or with MM_2.05_ fit separately from MM_0.94_+MM_3.0_ (MM_2+1_). Simulated-MM bases were scaled by a participant-specific MM_0.94/_MM_3.0_ amplitude-ratio estimated from 64-224 transients. Fit quality numbers (FQNs) between the five MM parameterization methods were calculated to compare fit quality. Intraclass and Pearson’s correlations were calculated to assess for agreement of GABA concentrations between the MM-suppressed GABA scan and the GABA+ scan with MMs fit out.

**Results:** GABA concentrations derived from fitting with simulated-/individual-MM bases and MM_2.05_ fit separately lead to good–excellent agreements and high correlations with MM-suppressed GABA concentrations. Fit quality was significantly (51–63%) worse when fit with the MM_together_ methods. GABA concentrations derived from fitting with a simulated MM basis demonstrated strong agreement with MM-suppressed GABA concentrations even when estimating the MM_0.94/_MM_3.0_ amplitude-ratio from a reduced number of transients (64–112).

**Conclusion:** A participant-specific MM basis with MM_2.05_ excluded/fit separately yielded accurate GABA concentrations and superior fits. An accurate estimation of the MM_0.94_/MM_3.0_ amplitude-ratio can be achieved from a short 3-minute MM scan.

## 1 Introduction

Gamma-aminobutyric acid (GABA) is the brain’s main inhibitory neurotransmitter and plays a significant role in a range of psychiatric, neurodegenerative, and neurometabolic diseases(1–4). As such, there is large interest in quantifying GABA non-invasively using in vivo proton magnetic resonance spectroscopy (^1^H-MRS). Due to substantial overlap in the ^1^H-MRS spectrum at 3T, GABA detection with conventional non-edited pulse sequences is difficult, as the GABA peaks at 1.9 ppm and 3.0 ppm are highly overlapped by the larger N-acetylaspartate (NAA) and creatine (Cr) peaks at 2.0 ppm and 3.02 ppm, respectively. As such, the robust detection of GABA with ^1^H-MRS requires specialized methods at 3T. Of these methods, J-difference editing (JDE) is the most commonly used and is considered the gold standard for GABA detection at 3T(5). JDE is typically performed using a MEGA-PRESS pulse sequence(6) which involves the acquisition of two sub-experiments where a selective 14 ms editing pulse alternates between being on- and off- resonance for GABA at 1.9 ppm (edit-ON) and ∼7.5 ppm (edit-OFF), respectively. Subtracting the two sub-acquisitions from one another (edit-ON – edit-OFF) yields a difference (DIFF) spectrum containing a GABA peak at 3.0 ppm.

While this approach removes the Cr signals overlapping GABA at 3.0 ppm, the editing pulses also partially invert macromolecule (MM) resonances at 1.72 ppm (MM_1.72_). As this leads to MM confounds in the edited 3.0 ppm peak, quantification of GABA from this spectrum is referred to as GABA+ (GABA + MM). Commonly, this MM confound is simply accepted(7–10). However, recent studies have demonstrated that accounting for the MM confound improves GABA measurements, where the pure GABA signal is a stronger indicator of behavior when compared to the GABA+ signal(11), and that MMs contributing to the GABA+ signal can skew GABA measurements in different brain regions(12).

To remove this confound, several methods such as inversion recovery(13) and MM-suppressed GABA editing(14) have been proposed. Inversion recovery-based methods exploit the substantial T1 relaxation differences between MMs and metabolites(15) to null the MM signal; however, these methods significantly reduce spectral signal-to-noise ratio (SNR) by 30–45% (assuming a MM T1 of 290-400 ms)(16) and increases T1-weighting in the spectrum(17). As such, symmetrical editing is more commonly used to detect MM-suppressed GABA. In this approach, the frequency of the edit-OFF pulse is shifted to 1.5 ppm so that it is symmetric about the MM_1.72_ resonance relative to the edit-ON pulse at 1.9 ppm(14). As the edit-ON and edit-OFF sub- acquisitions edit MM_1.72_ to the same degree, subtracting the two from one another results in a MM-suppressed DIFF spectrum.

This approach, however, requires a longer echo time (TE) of 80 ms relative to the more commonly used TE of 68 ms to allow for an increase in editing pulse duration from 14 ms to 20 ms to allow for greater frequency selectivity. This increased TE results in greater T2 relaxation effects and hence, a ∼20% reduction in the GABA SNR(18).

Furthermore, the symmetric positioning of the editing pulses about the MM_1.72_ resonance makes the sequence significantly more sensitive to frequency offsets originating from scanning after sequences with high gradient duty cycles (e.g. fMRI and DTI) and from participant movement, which is more pronounced in patient populations, thus leading to residual MM contamination(13). As such, several methods have been developed to prospectively correct for these frequency offsets. Notably, one method, metabolite cycling MEGA-PRESS (MC MEGA), uses a pre-inversion pulse in place of a conventional water suppression module that alternates between upfield and downfield inversion while leaving the water peak untouched(19). This allows for the correction of the transmit frequency at every transient without the need to acquire additional water references, and has shown to outperform other prospective frequency correction methods in suppressing co-edited MMs(19).

The MC MEGA sequence, however, is still vulnerable to large movements and does not completely eliminate the co-edited MM_3.0_ peak(20). This residual MM_3.0_ contribution to the 3.0 ppm peak can be even greater in brain regions with high B_0_ inhomogeneity due to the shifted transmit frequency of the editing pulse and broader distribution of spins across the frequency axis which break the symmetry of the editing pulses about MM_1.72_. Given the downsides to acquisition-based MM suppression methods, several modeling strategies to isolate GABA from MM have been proposed. Recently, it was shown that including a model of co-edited MMs as single Gaussian peaks at 0.94 ppm and 3.0 ppm to model MM_0.94_ and MM_3.0_, respectively, with a fixed 3/2 amplitude ratio led to improved fits and reduced residuals relative to fitting without a MM model(21). Another recent study demonstrated that using a MM basis derived from the cohort mean of experimental MM data improves fitting by allowing for more accurate modeling of the asymmetric MM peaks(22,23). Both fitting-based methods, however, assume a fixed MM_0.94_/MM_3.0_ amplitude ratio with negligible variations between participants. Literature, however, suggests that the MM profile can be significantly altered in different diseases such as multiple sclerosis and schizophrenia and can vary depending on the disease stage/state(24,25) and brain region of interest(26). The MM peak at 2.05 ppm (MM_2.05_) in the edited MM is also prone to contamination from residual metabolites and lipids, which can vary depending on movement, head shape, voxel positioning, and frequency offsets. Together, these technical and biological considerations can introduce variations into the MM profile, and neglecting to account for these variations could lead to inaccurate MM modeling and hence, inaccurate GABA concentration estimates.

As such, the goals of this study were to (1) assess whether the inclusion of a participant-specific MM basis improves the quantification of GABA from J-difference edited spectra and (2) evaluate the effects of different MM parameterization strategies on GABA quantification. JDE data targeting GABA were acquired with and without an inversion recovery pre-pulse to acquire metabolite-nulled MM-edited spectra and GABA-edited spectra, respectively, in healthy adults. Cohort mean, individual, and simulated MM bases were derived from the MM-edited data using two different fitting methods: one that included MM_3.0_, MM_2.05_, and MM_0.94_ altogether (MM_together_) and another that included MM_2.05_ as a separate basis function from MM_0.94_ and MM_3.0_ (MM_2+1_). The GABA+-edited spectra were then evaluated for fit quality, and their resulting GABA concentrations were evaluated for agreement with GABA concentrations derived from MM-suppressed GABA-edited scans.

## 2 Methods

### 2.1 Acquisition

All scans were performed on Philips Achieva 3T scanner with a 32-channel receive head coil. Thirteen healthy volunteers (9 female, 4 males; age 27 ± 7 years) gave informed written consent with local Institutional Review Board approval before any scan procedures were performed. Anatomical scans were acquired using a T1-weighted MP-RAGE(27) sequence with 1 mm^3^ isotropic resolution. All metabolite scans were performed using a MC MEGA sequence(19) which performs prospective frequency correction at every transient, making it ideal for minimizing MM contamination in MM-suppressed GABA-editing scans. All MRS data were acquired with second-order projection-based shimming, a repetition time (TR) of 2500 ms, 2000 Hz spectral width, and 2048 spectral points. MRS data were acquired in both the centrum semiovale (CSO), and the parietal lobe (Par), which are white matter (WM) and a gray matter (GM) majority regions, respectively. Voxel sizes were 50 x 20 x 20 mm^3^ in the CSO (13.3% GM), and 30 x 30 x 30 mm^3^ in the Par (57.7% GM) (Supplementary Table 1).

GABA+-edited spectra were acquired using a TE of 64 ms(18) with 15 ms sinc-Gaussian editing pulses and edit-ON/-OFF frequencies of 1.9 ppm and 7.5 ppm, respectively. For the Par voxel, additional MM-suppressed GABA-edited scans were acquired at a TE of 80 ms with 20 ms editing pulses and edit-ON/-OFF frequencies at 1.9 ppm and 1.5 ppm, respectively(28).

Metabolite-nulled, MM-edited scans were also acquired with a single-inversion recovery pre-pulse followed by a MEGA-PRESS sequence with identical TE, TR, and editing frequencies as the GABA+-edited scans. Other acquisition parameters included CHESS water suppression and an inversion time (TI) of 780 ms. This TI was selected based on the optimal nulling of NAA and GABA, which have similar T1 relaxation values at 3T (∼1320 ms)(29,30).

The minimum reporting standards for in vivo MRS(31) are further described in supplementary table 2.

### 2.2 MM Basis Formation

In the edited MM spectra, three MMs peaks were identified: MM_0.94_, MM_2.05_, and MM_3.0_. Estimating GABA free from MM in the GABA+-edited spectrum is made possible by the fully resolved MM_0.94_ peak which is not overlapped by signals from any confounding compounds. As spectral fitting software fits a uniform scaling factor to the MM basis, an accurate scaling factor for the entire MM basis, including MM_3.0_, can be estimated from the unambiguous MM_0.94_ peak. The remaining signal at 3.0 ppm can then be attributed to GABA as depicted in figure 1a. For the cohort mean and individual MM bases, the MM peaks were parametrized using a multi-Gaussian model with Gaussians centered at 0.05 ppm increments in regions-of-interests of 0.75–1.05 ppm (MM_0.94_), 1.90–2.25 ppm (MM_2.05_), and 2.85–3.20 ppm (MM_3.0_) to avoid fitting influence outside of the MM region-of-interest (Supplementary Figure 1). For the simulated MM basis, the MM_2.05_ peak was excluded from fitting, as the precise origin of these signals are unknown and thus cannot be accurately simulated(17), whereas the MM_0.94_ and MM_3.0_ signals are more established and can thus be modeled with density matrix simulations.

**Figure 1.**
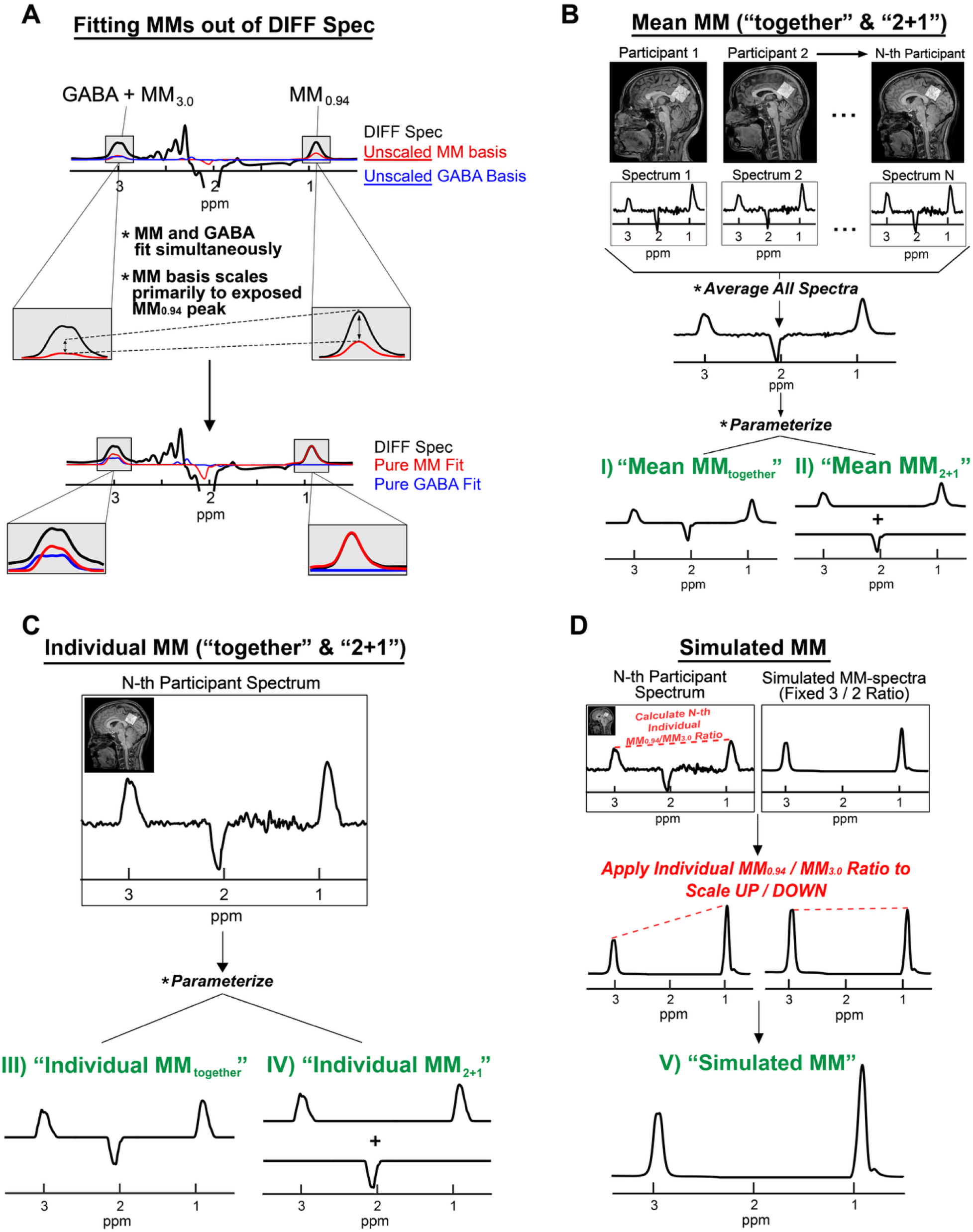
Schematic of fitting GABA and MMs in the GABA-edited spectrum and the different MM basis set generation methods. (A) The MM_0.94_ peak is fully resolved in th DIFF spectra and can therefore be used to scale the MM_3.0_ peak accordingly. This allows both the MM basis and GABA to be accurately fit. (B) The mean MM basis set was generated by averaging the metabolite-nulled DIFF spectra acquired from all participants in a given brain region. Each of the MM peaks in the resulting spectra were then parameterized with a multi-Gaussian model and recombined to form two types of basis sets: Mean MM_together_ with includes all three MM peaks in a single basis and Mean MM_2+1_ where MM_3.0_ and MM_0.94_ were included in separate basis from MM_2.05_. (C) Individual MM basis sets were constructed from each participant’s specific MM DIFF spectrum. Each of the MM peaks were parameterized using the same multi-Gaussian models in (B) and separated into Individual MM_together_ and Individual MM_2+1_ basis sets. (D) Combined simulated and individual MMs were generated by estimating the subject-specific MM_0.94_ / MM_3.0_ amplitude ratio and scaling the simulations of the MM_3.0_ peak relative to the MM_0.94_ peak.

The bases for the cohort mean and individual MMs were parameterized two different ways: (1) as a composite MM basis including all three MM peaks in a single basis (“together”) and (2) as two MM bases with MM_3.0_ and MM_0.94_ in one basis and MM_2.05_ in a separate basis (“2+1”). The latter parameterization was explored as the MM_2.05_ peak showed significant variability across participants likely due to lipid contamination and the presence of residual metabolites which could potentially affect the MM fit quality.

The “mean MM_together_” and “mean MM_2+1_” MM bases were constructed by parameterizing the cohort-averaged MM spectra for a given brain region using a multi-Gaussian model and finalized as either the “together” or “2+1” subcategories (Figure 1b). The “Individual MM_together_” and “Individual MM_2+1_” bases were constructed from each participant’s respective MM spectrum (Figure 1c). For the simulated MMs, the MM_3.0_ peak was modeled as Lys(17), and the MM_0.94_ peak was modeled as a combination of the BCAAs: leucine (Leu), isoleucine (Ile), and valine (Val)(17). The MM_0.94_/MM_3.0_ amplitude ratios were derived for each participant and used to scale the simulated MM peaks in the MM basis prior to fitting (Figure 1d).

### 2.3 Simulations

Density matrix simulations were performed using FID-A(32) with metabolite, Lys, and BCAA J-coupling constants and chemical shifts taken from prior literature(33–35). Simulations were performed over a 2-dimensional spatial plane split into 19 x 19 segments across the direction of the refocusing pulses. All metabolite simulations were apodized with a 3 Hz exponential line broadening filter.

### 2.4 Spectral Processing

The in vivo metabolite spectra were preprocessed using Gannet 3.1(36) with retrospective phase and frequency correction in the time domain(37), 3 Hz exponential line broadening, HLSVD water filtering(38), and zero-filling to 4096 points. The DIFF spectra were lipid filtered using a robust L2-regularization method(38). The spectral SNR values were estimated from the DIFF spectrum as the absolute maximum of the 2.0 ppm NAA peak divided by the standard deviation of the noise from 10–12 ppm. Linewidths were estimated from the DIFF spectrum as the full-width-at-half-max (FWHM) of the NAA peak at 2.0 ppm.

### 2.5 MRS Fitting

All GABA-edited spectra were fit using ProFit-1D(39) with basis sets containing GABA, glutamate (Glu), glutamine (Gln), glutathione (GSH), NAA, N-acetylaspartyl-glutamate (NAAG) and one of the given MM bases (see section 2.2). The simulated MM basis sets were simulated with a pool of Edit-ON frequency offsets ranging from -5 to +5 Hz. Each of the 320 acquired transients in the GABA+-edited scan had the frequency offset measured from the unsuppressed water peak to the nearest 0.5 Hz; the simulated basis applied that same corresponding frequency offset and was averaged across all transients(40). As the experimentally acquired MM bases included effects from both T2 relaxation and B_0_ inhomogeneity, Lorentzian and Gaussian line broadening terms were not included as part of the MM fitting process.

### 2.6 Analyses

As deriving participant-specific MM bases requires additional scans, we wanted to examine if significant reductions in MM scan time would still provide robust estimates of the MM_0.94_/MM_3.0_ amplitude ratio. MM_0.94_/MM_3.0_ amplitude ratios were extracted with 224 number of signal averages (NSA) (full NSA; ∼9.3 min. scan), 112 NSA (∼4.6 min scan), 80 NSA (∼3.3 min scan), and 64 NSA (∼2.6 min scan) and used to scale the given simulated MM basis. Fit quality numbers (FQNs) were calculated in Profit-1D, as previously described(39,41), to assess the fit quality in the 0.6–4.05 ppm range but excluding the 1.15–1.75 ppm range to avoid confounds from lipids. In vivo GABA concentrations were estimated using each MM modeling strategy and evaluated for agreement with the MM-suppressed GABA concentrations. GABA concentrations across each MM modeling strategy were compared between the Par and CSO.

### 2.7 Statistical Analyses

Pearson’s correlation coefficient and two-way mixed effects intraclass correlation coefficients, denoted as ICC(3,1), where 3 indicates a two-way mixed effects model and 1 indicates the single measurement reliability, were calculated to assess the agreement between GABA concentrations estimated from the different MM modeling strategies and the MM-suppressed GABA scans. ICCs were classified as follows: poor (ICC < 0.4), moderate (0.4 ≤ ICC < 0.59), good (0.6 ≤ ICC < 0.74), and excellent (ICC ≥ 0.75)(42). All statistical analyses were conducted in MATLAB. Outliers were identified as > 3 scaled median absolute deviations from the median and excluded from statistical testing. T-tests were used to assess statistically significant differences in FQNs and GABA levels across modeling strategies.

## 3 Results

Figure 2 shows the SUM (Figure 2a) and DIFF (Figure 2b) spectra taken from an inversion recovery sequence performed at TIs ranging from 720–840 ms. Although the three large singlets, namely tCr (phosphocreatine + creatine), tCho (phosphocholine + glycerophosphocholine + choline), and tNAA (NAA + NAAG) are optimally nulled at different TIs, a TI of 780 ms optimally nulled the tNAA singlet at 2.0 ppm (Figure 2a — shaded). This TI was chosen for subsequent MM basis acquisitions since tNAA, unlike other large singlets in the spectrum, would not be subtracted out in the DIFF spectrum and could confound measurement of the nearby MM_2.05_ resonance (Figure 2b).

**Figure 2.**
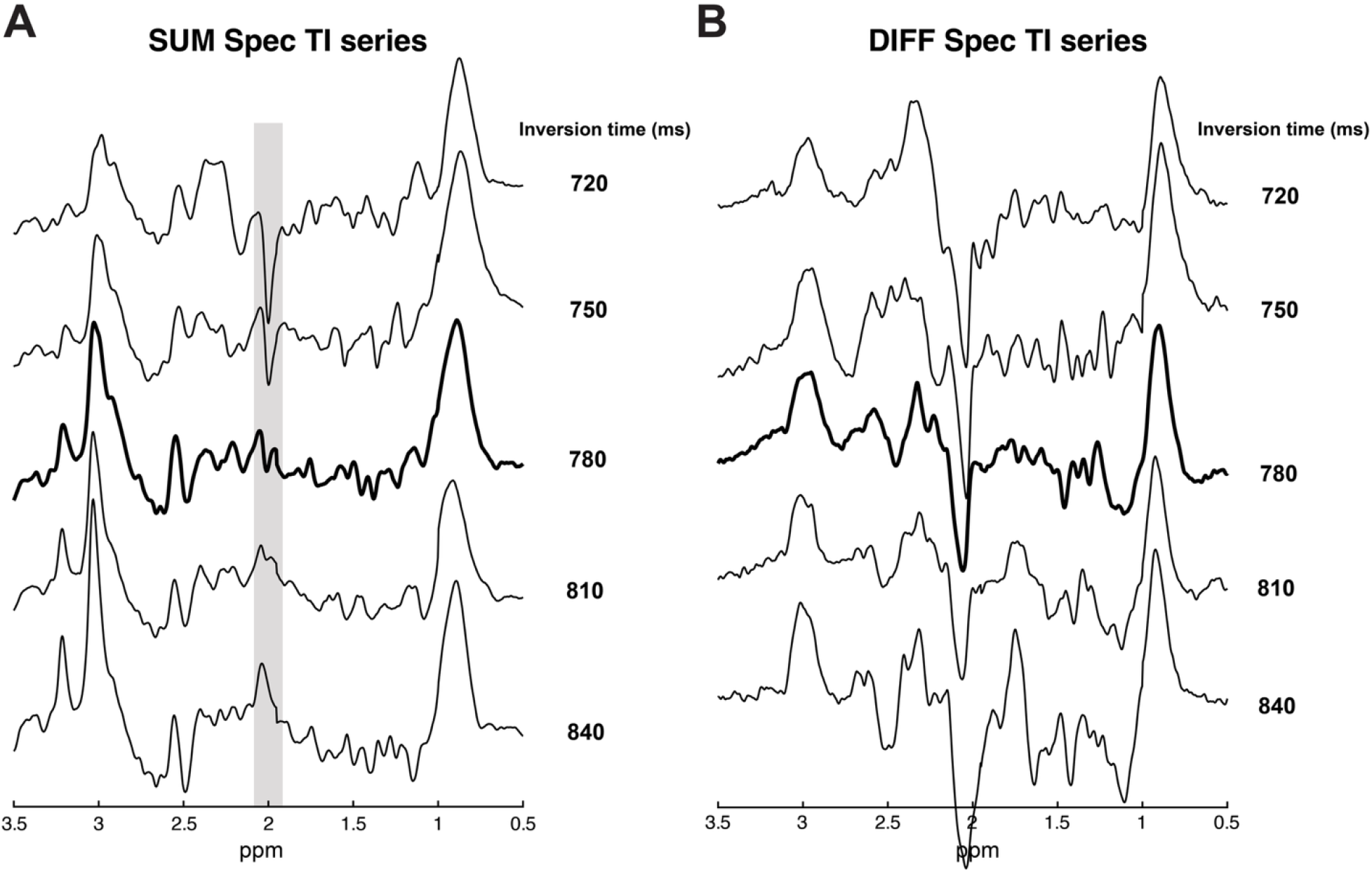
Example spectra acquired with an inversion-recovery sequence performed at inversion times of 720 ms to 840 ms at 30 ms increments in one participant. Both the SUM (A) and DIFF (B) spectra are shown for each TI. The spectra acquired at a TI of 780 ms are bolded to emphasize optimal metabolite nulling for tNAA (shaded region) at the expense of residual tCr and tCho, as these peaks are subtracted out in the difference spectra.

Additionally, tNAA has a T1 relaxation time similar to GABA at 3T; as such, GABA would be optimally nulled at a similar TI (29,30).

Figure 3a shows example voxel placements in the GM-heavy Par (left) and WM-heavy CSO (right) while figures 3b and 3c show the mean ± standard deviation of the MM-edited spectra and GABA-edited spectra, respectively. Both the MM and GABA-edited spectra were of excellent quality, with SNR values of 111–165 and mean tNAA FWHM values of 4.83–5.38 Hz which are considered excellent per experts’ consensus(43) (Table 1).

**Figure 3.**
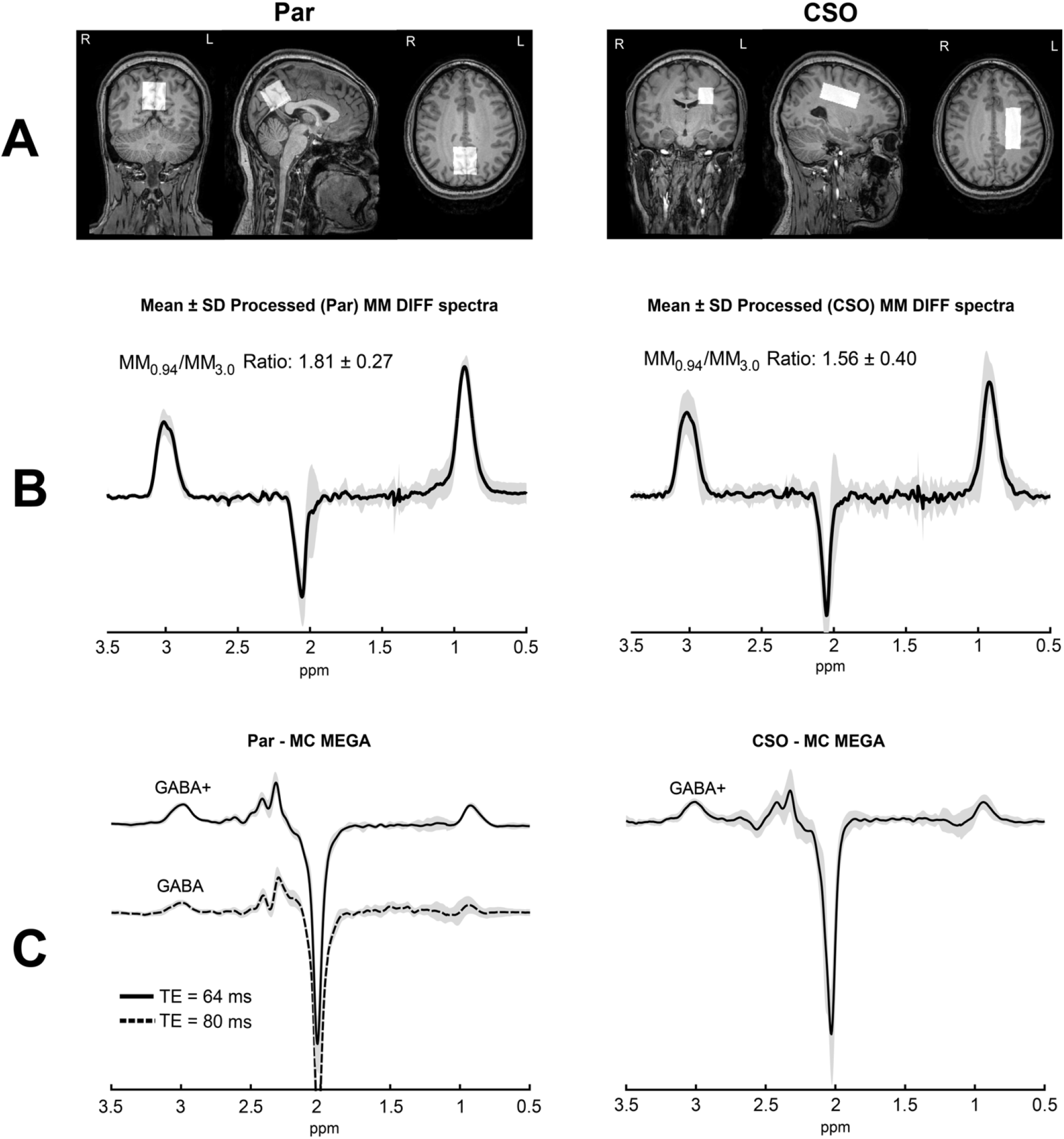
Example voxel placements (A), metabolite-nulled MM-edited spectra (B), and GABA-edited spectra (C). (A) Representative voxel placements in the parietal lobe (30 x 30 x 30 mm^3^) and centrum semiovale (50 x 20 x 20 mm^3^). (B) Mean ± standard deviation of the metabolite-nulled spectra across all participants in the Par (left) and CSO (right), where the MM spectra display a higher MM_0.94_/MM_3.0_ amplitude ratio in the Par than the CSO. (C) Mean ± standard deviation of the GABA-edited spectra acquired from the Par (left) and CSO (right). All GABA-edited spectra (C) are of high quality, with an observable decrease in the GABA 3.0 ppm amplitude in the MM-suppressed spectra.

**Table 1.**
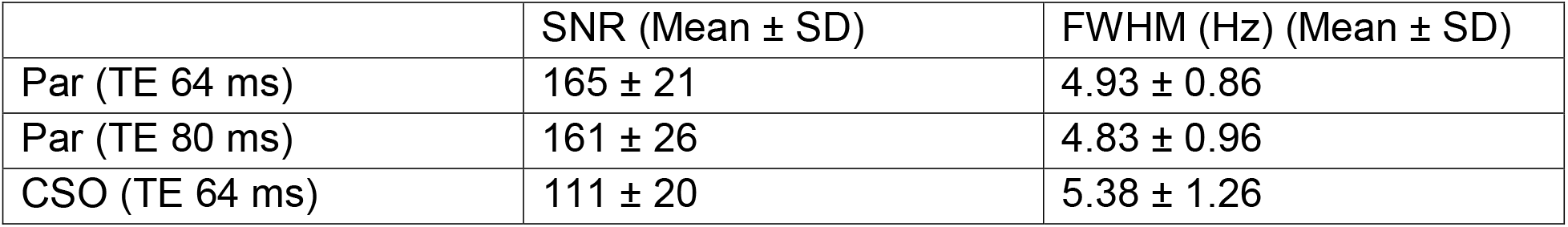
SNR and FWHM values for the in vivo metabolite spectra.

| | SNR (Mean ± SD) | FWHM (Hz) (Mean ± SD) |
| --- | --- | --- |
| Par (TE 64 ms) | 165 ± 21 | 4.93 ± 0.86 |
| Par (TE 80 ms) | 161 ± 26 | 4.83 ± 0.96 |
| CSO (TE 64 ms) | 111 ± 20 | 5.38 ± 1.26 |

The mean ± standard deviation of the MM_0.94_/MM_3.0_ peak amplitude ratio displayed a trend (p = 0.096) of being higher in GM vs. WM with values of 1.81 ± 0.27 and 1.56 ± 0.40, respectively, which is consistent with prior literature(44). While the MM-edited spectra are highly consistent between participants, some variation can be seen near the MM_2.05_ resonance, which is most likely a result of contamination from lipid and residual tNAA (Figure 3b). The GABA-edited spectra were shown to be highly consistent between participants, particularly in the 3.0 ppm region where GABA is located. In the MM-suppressed scans, the 3.0 ppm region is substantially smaller, indicating effective MM suppression for all participants. In these scans, MM contamination was also minimal, with a median absolute water frequency shift of 0.41 Hz across all MM-suppressed scans.

Figure 4 shows linear regression plots of the MM-suppressed GABA concentrations (x-axis) compared to the simulated MM, mean MM_together_, mean MM_2+1_, individual MM_together_, and individual MM_2+1_ fitted GABA concentrations (y-axis). ICC(3,1), r-values, and corresponding p-values for each modeling strategy are expressed on each plot. Among the five modeling strategies, GABA concentrations estimated from the simulated MMs (Figure 4a) and individual MM_2+1_ (Figure 4e) displayed the best agreement with those estimated from the MM-suppressed scans. This agreement was reflected in their ICC(3,1) scores of 0.68 (good) and 0.90 (excellent) for simulated MMs (Figure 4a) and individual MM_2+1_ (Figure 4e), respectively. GABA concentrations estimated from both the simulated MMs and individual MM_2+1_ modeling strategies were also strongly correlated with those estimated from the MM-suppressed scans with statistically significant correlations of r = 0.6844 and r = 0.9109 (p < 0.05), respectively. In contrast, GABA concentrations estimated from the mean MM_together_, mean MM_2+1_, and individual MM_together_ modeling strategies showed poor agreement with those from the MM-suppressed scans with low ICC(3,1) < 0.4, and correlations of r = 0.18–0.37 (Figure 4b–d), none of which were statistically significant.

**Figure 4.**
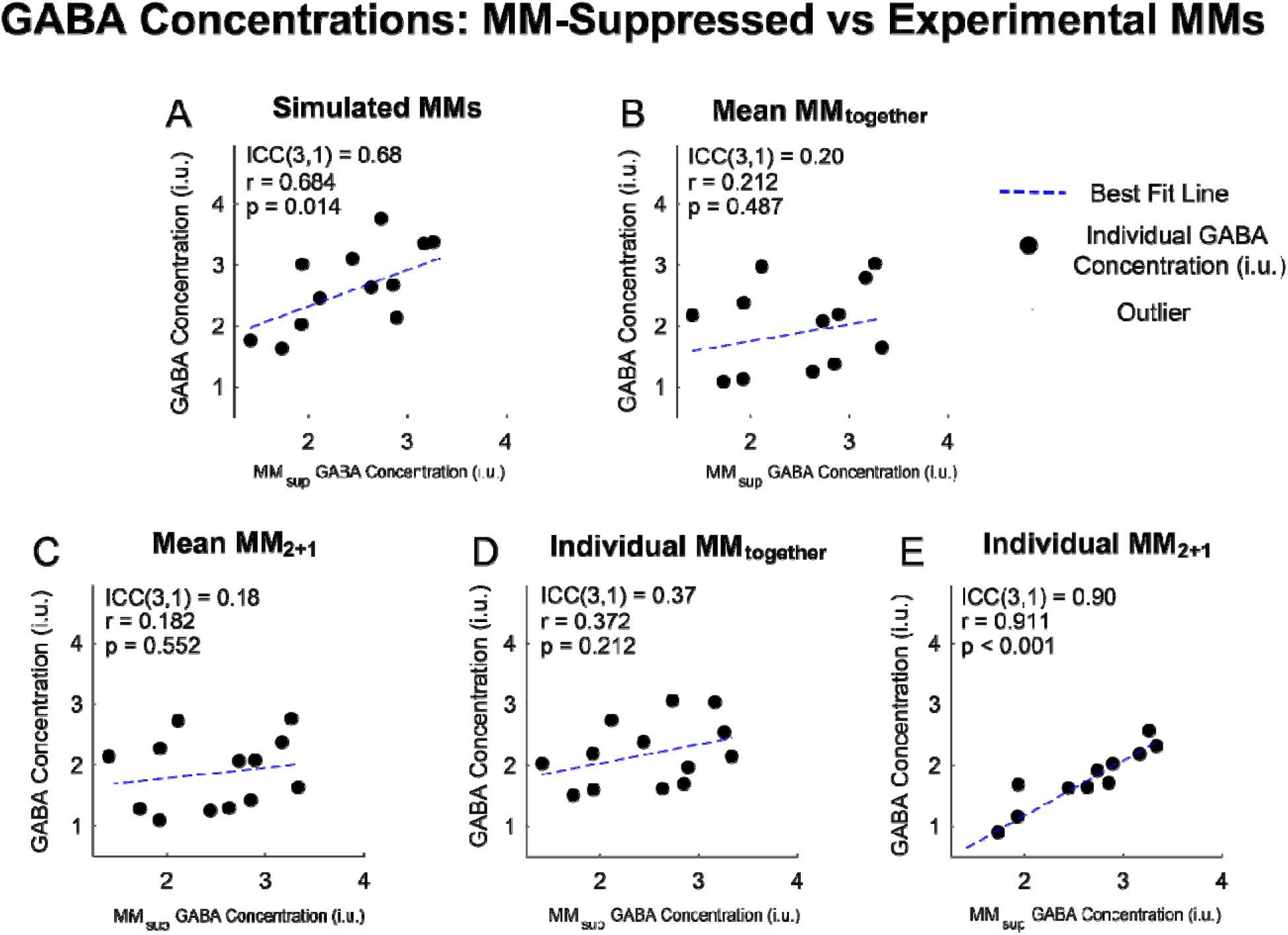
Linear regression plots between the GABA levels estimated from modeling the MM in the GABA+ acquisitions vs. the GABA levels estimated from the MM-suppressed (MM_sup_) scan. Two-way mixed effects intraclass correlations (ICC (3,1)) and Pearson’s correlation coefficients are also shown for each method. Only the simulated MM (A) and Individual MM_2+1_ (E) methods resulted in high ICC(3,1) and r-values indicating an agreement with GABA concentrations reported from MM-suppressed scans. GABA concentrations estimated from fitting with the cohort mean MMs (B and C) did not show significant correlations with GABA concentrations estimated from the MM-suppressed scans. Outliers are marked as red crosses.

Differences in fit quality between the modeling strategies were also assessed from the FQN values. In both regions, the FQN values using the simulated MM strategy were significantly lower than those using the Mean MM_together_ and Individual MM_together_ bases by 51–63% (p < 0.001) and 62–63% (p < 0.05) in the Par (Figure 5a) and CSO (Figure 5b), respectively. Fitting with mean MM_2+1_ and individual MM_2+1_ did not lead to statistically significant increases in FQN relative to the simulated MMs. For both the mean and individual MM fitting strategies, fitting with the “MM_2+1_” method opposed to “MM_togther_” resulted in 23–38% and 32–33% lower median FQNs in the Par and CSO, respectively, though this finding was not statistically significant (Figure 5). GABA concentrations were higher in the Par than the CSO across all fitting methods, although this finding was only statistically significant when fitting with simulated MMs (p = 0.034) (Supplementary Figure 2).

**Figure 5.**
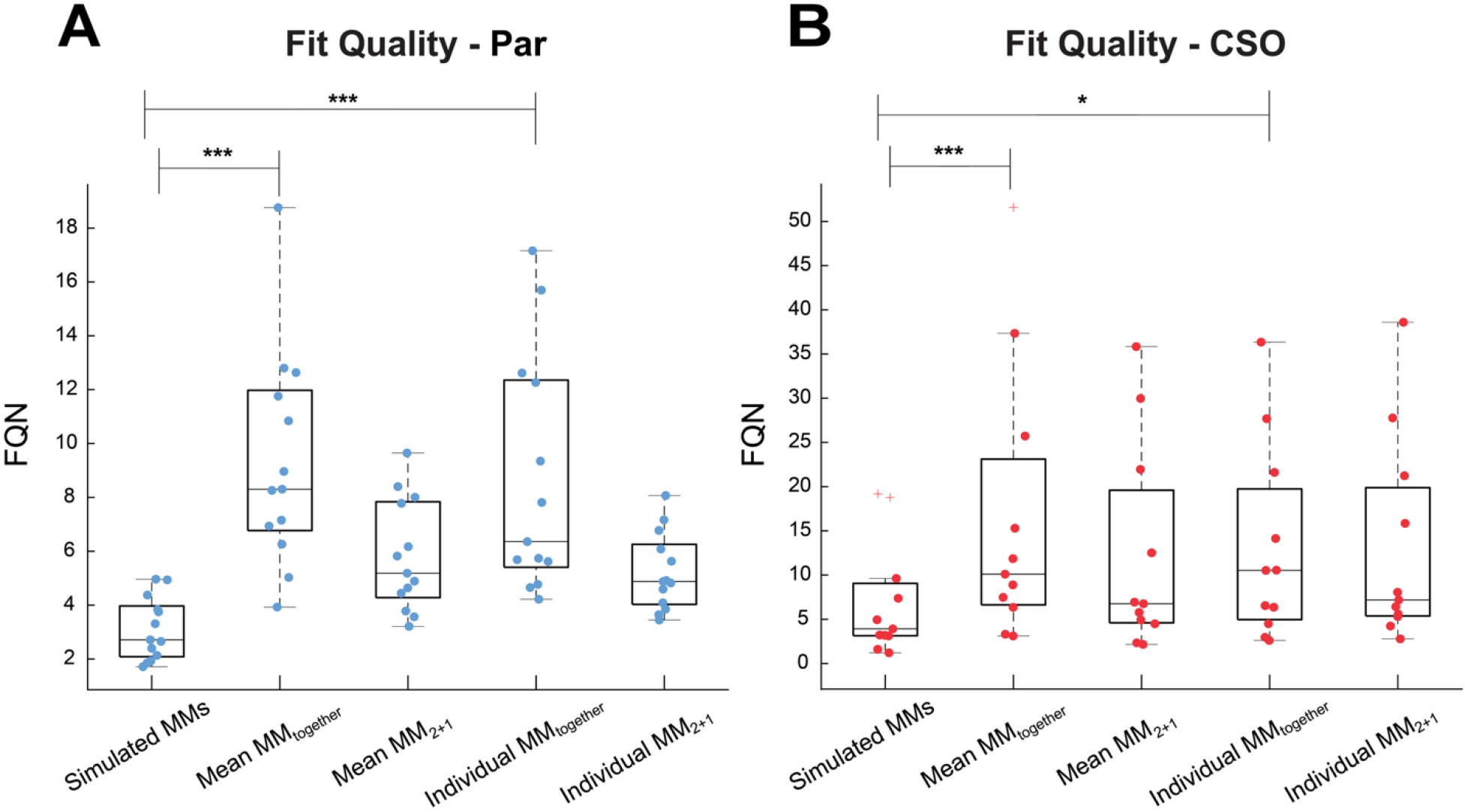
Fit quality numbers (FQNs) for the different modeling strategies in the (A) Par and (B) CSO regions. FQN values were calculated in the 0.6 – 4.05 ppm range, excluding the 1.15 – 1.75 ppm range which contained lipid resonances that were not included in the model. Fitting with simulated MMs resulted in the lowest FQNs. Fitting strategies that included the MM_2.05_ peak as part of a composite basis function (Mean MM_together_ and Individual MM_together_) also resulted in significantly increased FQN values. *P≤0.05, *P≤0.01, ***P≤0.001. Outliers are marked as red crosses.

While acquiring a participant-specific MM spectrum substantially increases the total scan session duration, obtaining an accurate estimate of the MM_0.94_/MM_3.0_ amplitude ratio may still be achievable even when using a greatly reduced NSA, and hence, a reduced scan time. As such, we examined the effect of NSA on the MM_0.94_/MM_3.0_ amplitude ratio estimation and its impact on the GABA levels when fitting with the simulated MM strategy. Figure 6a shows the MM_0.94_/MM_3.0_ amplitude ratio as a function of NSA included in the MM-edited spectral reconstruction. Initially, the estimated MM_0.94_/MM_3.0_ amplitude ratios vary substantially as a function of NSA but begin to even out after ∼60 transients, though there is still a relatively wide distribution of MM_0.94_/MM_3.0_ ratios even after plateauing. The effect of NSA is shown further in figure 6b which shows the percent difference in the MM_0.94_/MM_3.0_ amplitude ratio relative to the participant-specific MM_0.94_/MM_3.0_ amplitude ratio estimated from the MM spectrum when reconstructed with all acquired transients. It can be seen here that the percent difference in the estimated MM_0.94_/MM_3.0_ amplitude ratio steeply decreases and starts to plateau at ∼60 NSA (∼27% of the total NSA of 224) at a ∼15% difference. Furthermore, when fitting with simulated basis sets scaled by the MM_0.94_/MM_3.0_ amplitude ratios estimated with lower NSA values of 112, 80, and 64, the ICC(3,1) values did not change substantially and remained “good” at 0.59 to 0.69, which are all similar to that of the fully acquired NSA (ICC(3,1) = 0.68) (Figure 6c–f). Correlations between the fitted GABA concentrations and MM-suppressed GABA concentrations also remained strong with r-values of 0.61 – 0.7 (p < 0.05) (Figure 6c–f). There were also no significant changes in the median (interquartile range) of GABA concentrations when fitting with amplitude ratios extracted with 224, 112, and 64 NSA, with values of 2.89 (1.44), 2.78 (1.51), and 2.81 (1.38) institutional units (i.u.), respectively (p > 0.15).

**Figure 6.**
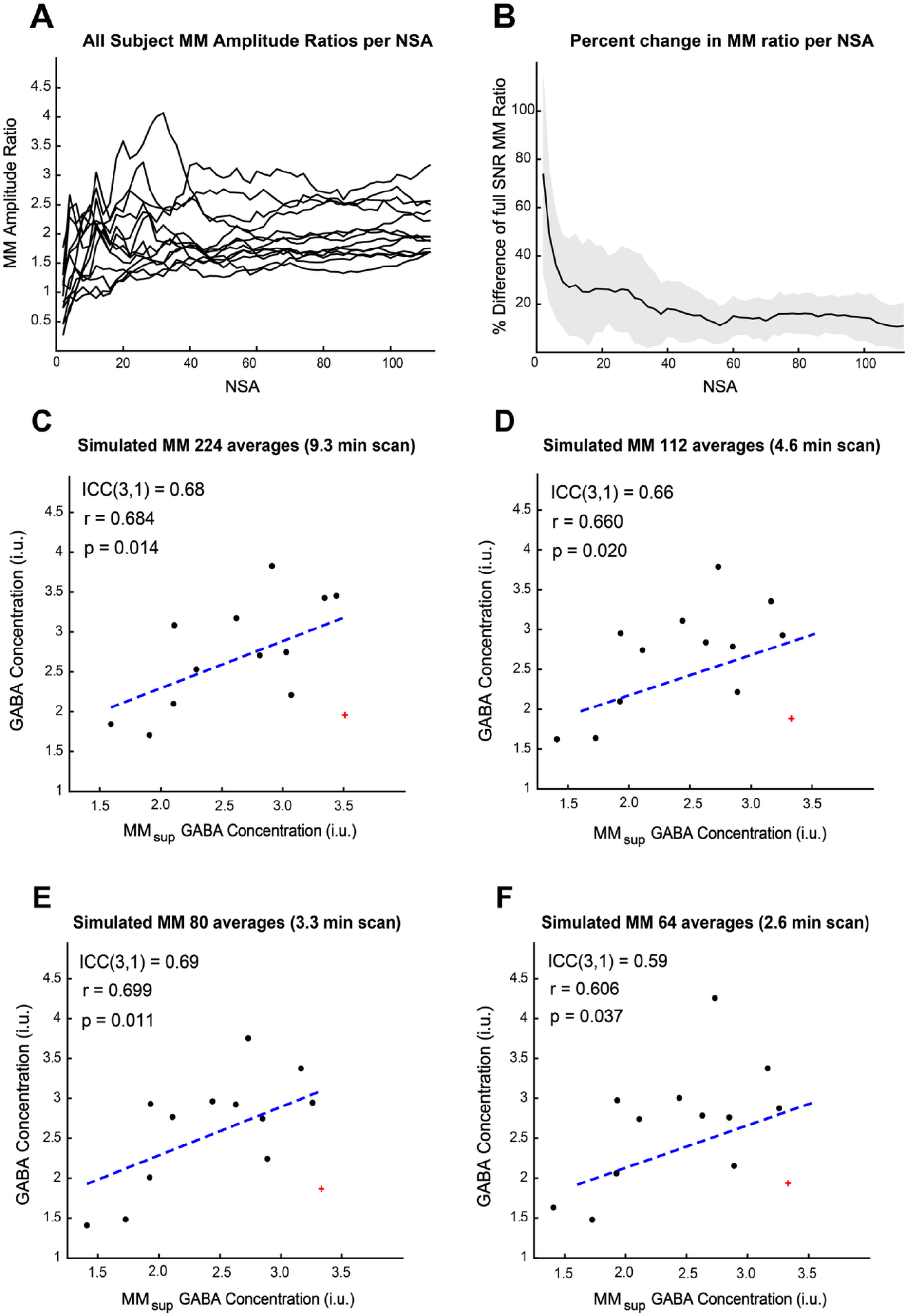
Influence of the number of signal averages (NSAs) on the estimated MM_0.94_/MM_3.0_ amplitude ratios (A) and (B) and agreement with the MM-suppressed GABA concentrations (C–E). (A) Participant-specific MM_0.94_ / MM_3.0_ ratios as a function of NSA. (B) Participant-specific MM_0.94_ / MM_3.0_ ratios from (A) expressed as a percentage relative to MM_0.94_ / MM_3.0_ ratio measured from MM spectra reconstructed from all 224 NSAs. An observable plateau in estimated MM_0.94_ / MM_3.0_ ratios can be seen at ∼64 NSA. (C–F) ICC(3,1) plots comparing GABA concentrations derived from fitting with a simulated MM basis and amplitude ratios derived from a scan with (C) 224 NSA, (D) 112 NSA, (E) 80 NSA, and (F) 64 NSA compared to GABA concentrations estimated from the MM-suppressed (MM_sup_) scans. Corresponding correlation and p-values are also displayed on each subfigure. All NSA values examined still possess high correlation coefficients. Outliers are marked as red crosses.

## 4 Discussion

In this study, five MM modeling strategies to estimate GABA separate from the MM in GABA+ spectra were investigated. Overall, it was found that fitting with a MM basis estimated from experimentally-derived, participant-specific information may provide better GABA estimates than fitting with a cohort mean MM basis. Specifically, GABA concentrations estimated from fitting the GABA+ spectra with MM basis sets derived from participant-specific information showed good to excellent agreement with the MM-suppressed GABA concentrations with high ICC(3,1) and Pearson’s correlation coefficients. In the case of the individual MMs, this finding was found to be true only when MM_2.05_ was fit separately (individual MM_2+1_).

Recent work has shown that the inclusion of an experimentally-derived cohort-averaged MM basis set improves GABA fitting(23). This same study, however, did not evaluate the effects of using a participant-specific MM basis, include a comparison to MM-suppressed GABA-editing scans, or systematically investigate different MM parameterization strategies. A prior study on short TE PRESS data, however, suggested that fitting with an experimentally-derived participant-specific MM basis does not confer any additional benefit in comparison to a cohort mean MM basis(45). In contrast, this study found that including participant-specific information in the MM basis allows for more accurate GABA estimates when measured with J-difference editing.

This discrepancy may be due to the lower metabolite signal intensities relative to the MM in the J-difference edited spectra, resulting in GABA estimates that are more sensitive to inter-participant MM variability. Furthermore, it is possible that the MMs measured from the GABA+-edited spectra may vary more between participants in comparison to other MMs.

### 4.1 MM_2.05_

The amino acids underlying the MM_2.05_ resonance have not been comprehensively characterized(17). As such, MM_2.05_ was excluded from the simulated MM basis, as inaccurate modeling of this peak could introduce error. Unlike the other MMs in the acquired MM spectra, the MM_2.05_ peak varied significantly between participants; this is likely due to residual tNAA contamination and lipid contamination originating from the skull, as this region is not shimmed(46), and is partially inverted by the 1.9 ppm editing pulse, thus contaminating the DIFF spectra. Considering that the MM_2.05_ peak showed substantial variation between participants, we examined the effect of fitting MM_2.05_ separately from the MM_3.0_ and MM_0.94_ basis (MM_2+1_). Here, it was found that the fit quality improved when MM_2.05_ was fit separately with both the cohort mean MM and individual MM fitting strategies. As the MM_together_ fitting-based strategies adjust the amplitudes to all three MM peaks together, these results suggest that the inclusion of a poorly defined MM_2.05_ peak negatively impacts the scaling of the MM_0.94_ and MM_3.0_ peaks, and hence the GABA level estimates.

### 4.2 MM-Fitting vs MM-suppressed GABA editing

MM-suppressed GABA editing has shown promise for obtaining more specific GABA measurements(1,47,48); however, this method still has several technical downsides. Prior simulations have indicated that MM-suppressed GABA editing does not fully suppress the co-edited MM signal, and roughly 10–15% of the “pure GABA” signal can be attributed to residual MM_3.0_(20). This method also requires a significantly longer TE which decreases GABA SNR(18) by ∼20% due to GABA’s short T2 relaxation time of 88 ms(49). The increase in TE also eliminates the opportunity for the simultaneous detection of other inhibitory metabolites such as glycine, which is possible when using a shorter TE of 64 ms(18). Another drawback of MM-suppressed GABA editing is the reliance of the symmetrical positioning of the editing pulses about the co-edited MM_1.72_ resonance. This requirement decreases the method’s robustness, particularly in patient populations where substantial movement can lead to variable contributions from co-edited MMs, thereby decreasing the reliability of GABA estimates. This effect is especially problematic when acquiring data from brain regions with significant B_0_ inhomogeneity such as the prefrontal cortex and hippocampus, which are regions of great interest for studying GABA in psychiatric pathologies such as mania(50), social anxiety disorder(51), and schizophrenia(52,53). In these regions, the MM_1.72_ peak becomes substantially broader as the spins become more widely distributed across the frequency axis. This breaks the symmetrical alignment of the edit-ON and OFF pulses and results in unwanted MM contribution to the 3.0 ppm GABA peak even in the MM-“suppressed” spectrum.

While MM-suppressed GABA editing comes with several downsides, these limitations were mitigated here with several study design choices. Firstly, the brain regions of interest are located far away from any air-tissue interfaces that might cause susceptibility effects and thus have favorable shimming conditions. Secondly, all MRS scans were performed in cohort of young healthy participants, who tend to be more MR-compliant relative to patient and older populations. Thirdly, MC MEGA was used to acquire the GABA-edited data (19); this sequence enables the simultaneous acquisition of the unsuppressed water peak and metabolite data at every transient, thus allowing for prospective frequency correction at every transient without a reduction in scan time. As a result, MC MEGA was found to increase the robustness of MM-suppressed GABA-editing to frequency offsets(19). Similar to what was found previously, we observed low absolute water frequency offsets, indicating minimal contamination from MMs.

Considering these favorable scan conditions, the GABA levels estimated here from the MM-suppressed scans provide a good reference for the GABA levels estimated from fitting the GABA+ spectra with the different MM modeling strategies.

### 4.3 Simulated vs acquired MM basis

In this study, high-quality MM-edited spectra were acquired with a relatively long scan time of 9.3 minutes, which can be prohibitively long for studies that need to examine multiple regions of interests or require additional information from other MRI scans. It should be noted, however, that both the fully acquired MM bases and simulated MM bases performed comparatively well at estimating GABA concentrations even though the multiplet patterns of the MMs were not fully modeled by the simulations. This is likely due to inaccuracies in modeling the MM spin systems, which have yet to be fully characterized in the human brain.

A major benefit of the simulated MM approach, however, is that it allows for accurate GABA measures as long as sufficiently accurate estimations of the MM_0.94_/MM_3.0_ amplitude ratio are obtained. As seen here, this can be done in the Par with just 64–80 NSA which corresponds to a 2.6–3.3 minute scan, 2.8x–3.5x faster than what was originally acquired. The MM_0.94_/MM_3.0_ amplitude ratios estimated from this NSA range resulted in GABA concentrations that maintained a good to moderate agreement with MM-suppressed scans, similar to what was found for the full NSA data. As such, it may be more time efficient to acquire an abbreviated MM-edited scan to estimate the participant-specific MM_0.94_/MM_3.0_ amplitude ratio and incorporate it into the simulated MM basis to estimate GABA free of MM.

### 4.4 Edited MM in the Par and CSO

It should be noted that lower FQN values, indicating higher fit qualities, were found in the GM-majority Par region than in the WM-heavy CSO across all modeling strategies. Between the five modeling strategies, the individual MM_2+1_ basis was the only fitting strategy that led to consistent estimations of GABA in the CSO. This is likely due to the lower spectral SNR in the CSO due to the region’s larger distance from the receive coils and lower overall MM and GABA levels, which have been shown to be∼28% and 50% lower, respectively, in WM than in GM(54,55). As such, a participant-specific experimentally determined MM basis may not be suitable in instances where WM-heavy regions or any other low-SNR regions are being scanned due to the time cost of acquiring a MM spectrum with sufficiently-high SNR. In such cases, MM-suppressed GABA-editing scans with prospective frequency correction (e.g. MC-MEGA) may be better suited.

## 5 Conclusion

In summary, five different strategies to model MMs out of the GABA+ signal were compared. It was found that fitting strategies that incorporated a participant-specific component in the MM basis outperformed cohort mean MM bases. Additionally, fitting strategies that either excluded or fit MM_2.05_ independently from the other MMs demonstrated superior performance relative to those that included MM_2.05_ in the same basis as the other MMs. Simulating MM bases and scaling the MM_0.94_ and MM_3.0_ peaks by amplitude ratios obtained from participant-specific MM scans with a reduced number of signal averages was found to show promise in providing MM-free GABA values without substantially increasing scan time. Taken together, these participant-specific MM modeling strategies may be preferable when scanning in brain regions with higher B_0_ inhomogeneity and in patient populations that are more prone to movement.

## Acknowledgements

This work was supported by the National Cancer Institute (P30CA142543) and American Cancer Society (IRG-24-1322075-19-IRG)

