## Supporting Information for "Participant-specific macromolecule information improves quantification of GABA-edited spectra at 3T"

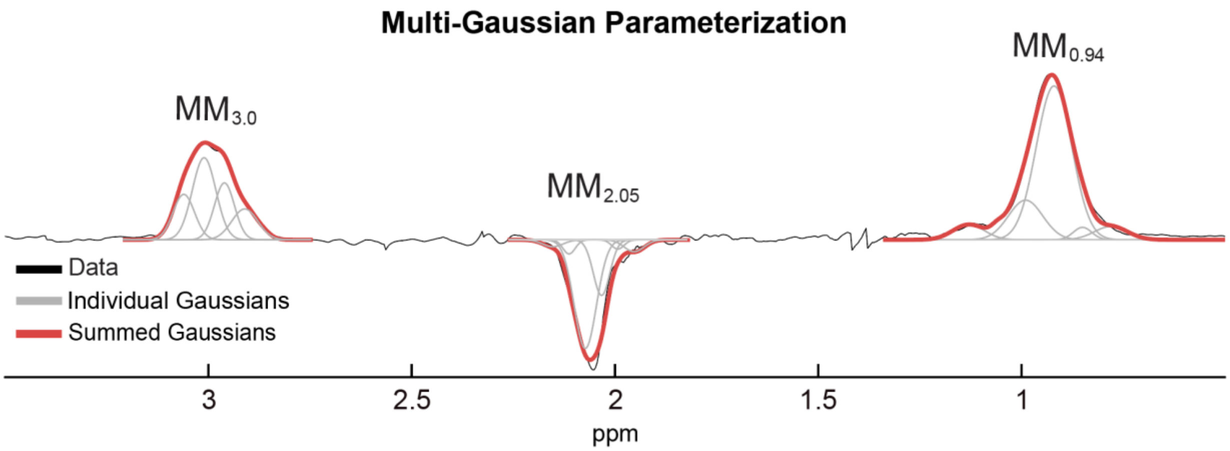


S Fig 1. Example multi-Gaussian parameterization in a healthy participant. All acquired MM spectra were parametrized using a multi-Gaussian model which consists of Gaussian peaks centered at 0.05 ppm increments within the 0.75–1.05 ppm, 1.90–2.25 ppm, and 2.85–3.20 ppm regions-of-interest and fit with a non-linear least squares optimization. All regions outside of the region-of-interest were set to zero to avoid signals outside the MMs of interest that may complicate fitting.

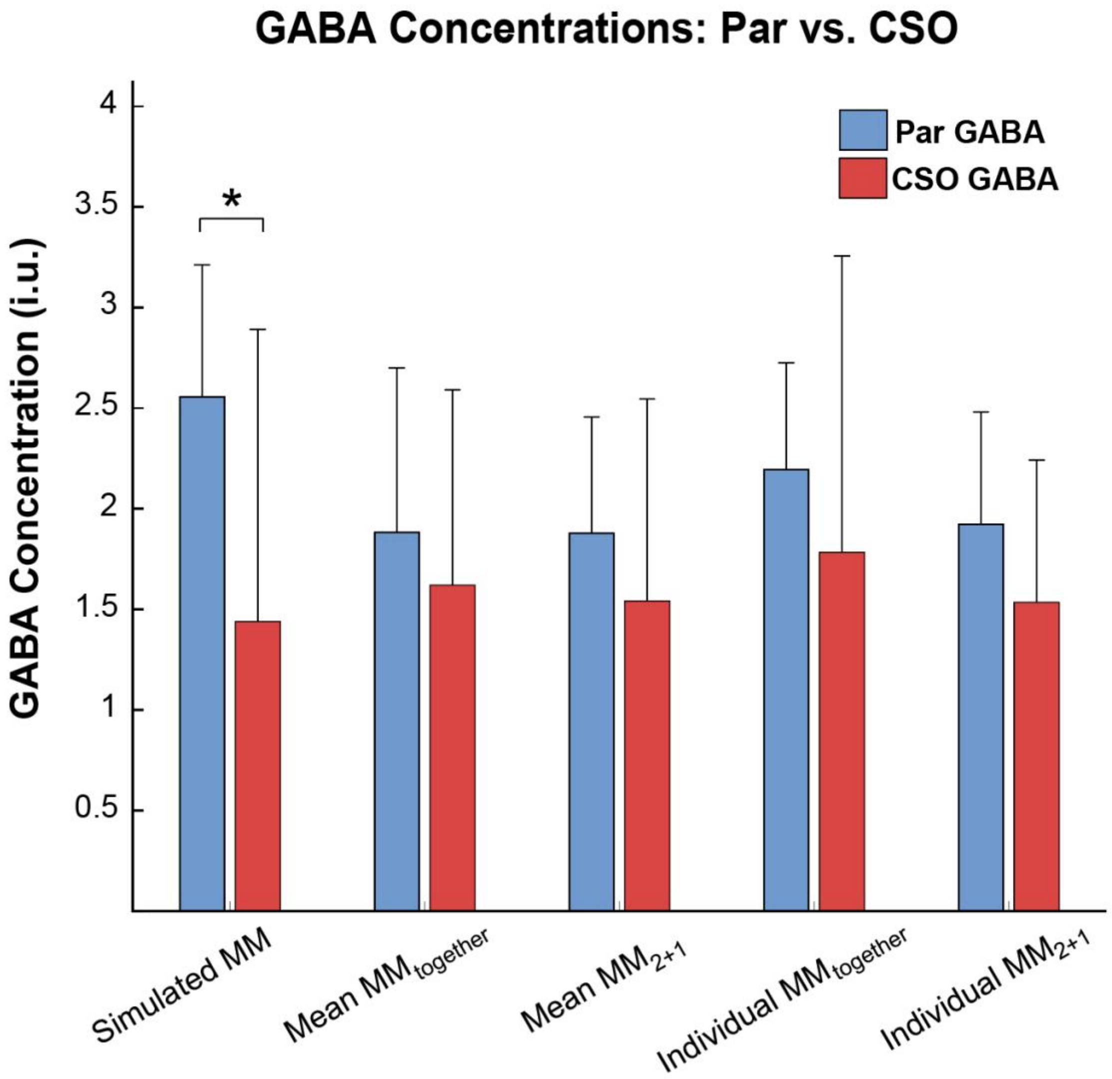


S Fig 2. GABA concentrations in the Par and CSO. Bar plots with mean and SD of the GABA concentrations in the Par and CSO with all fitting methods. GABA in the Par displayed a trend of being higher for all fitting strategies but was only statistically significant when fitting with the simulated MM basis. *P≤0.05.

Supplementary Table 1. Gray matter and white matter fractions for all voxel locations scanned.

| | Gray Matter % (Mean ± SD) | White Matter % (Mean ± SD) |
|---|---|---|
| Par | 57.7 ± 3.6% | 32.4 ± 3.9% |
| CSO | 13.3 ± 4.3% | 85.0 ± 5.1% |

Supplementary Table 2. Minimum Reporting Standards in Magnetic Resonance Spectroscopy checklist

Hardware

| | |
|---|---|
| Field Strength | 3T |
| Manufacturer | Philips |
| Model (Software) | Achieva, software release 5.7.1 |
| RF Coil | 32-channel $^{1}$H head coil |
| Additional Hardware | N/A |

Acquisition

| | |
|---|---|
| Pulse Sequence | MC-MEGA; MEGA-PRESS |
| Volume of Interest (VOI) | Par; CSO |
| Nominal VOI size | Par 27 mL; CSO 20 mL |
| RF Coil | 32-channel head coil |
| Repetition Time (TR) and Echo Time (TE) | TR = 2.5 s and TE = 64 and 80 ms |
| Number of averaged spectra | 320 averages; 224 averages |
| Additional Parameters | 2000 Hz spectral width, 2048 spectral points |
| Water Suppression Method | Metabolite Cycling / Philipps' CHESS |
| Shimming Method | Pencil beam (PB)-auto, 2nd order |
| Triggering or Motion Correction | N/A |

Analysis and Data Quality

| | |
|---|---|
| Analysis Software | Gannet 3.1, Profit-1D |
| Processing steps deviating from reference | N/A |
| Output Measure | Institutional Units |

| | |
|---|---|
| Quantification Reference | Unsuppressed water |
| Reported Variables | SNR (based off the NAA peak) and full-width-at-half-max of the NAA peak both derived from the DIFF spectra |
| Data exclusion criteria | Visual inspection (lack of interpretable spectrum, movement) |
| Sample spectrum | Figure 3 |
